# Benchmarking generative scaffold design methods for peptide engineering in TCR–MHC complexes

**DOI:** 10.64898/2026.01.22.701133

**Authors:** Linhui Xie, Gia-Bao Dam, Yashvi Patel, Lilian Denzler, Yanjun Shao, Ruimin Wang, Etienne Caron, Yoshiaki Yasumizu, David A. Hafler, María Rodríguez Martínez

**Affiliations:** Department of Biomedical Informatics and Data Science, Yale School of Medicine, 101 College Street, New Haven, 06510, CT, USA; Department of Statistics and Data Science, Yale University, 219 Prospect Street, New Haven, 06511, CT, USA; Department of Physics, Yale University, 217 Prospect Street, New Haven, 06511, CT, USA; Department of Structural and Molecular Biology, University College London, Darwin Building, Gower Street, London, WC1E 6BT, UK; Department of Immunobiology, Yale School of Medicine, 300 George Street, New Haven, 06511, CT, USA; Department of Neurology, Yale School of Medicine, 300 George Street, New Haven, 06511, CT, USA

**Keywords:** TCR-pMHC, *de novo* peptide design, RFdiffusion, protein diffusion models, computational immunology, benchmark

## Abstract

De novo peptide design at T cell receptor–peptide–major histocompatibility complex (TCR–pMHC) interfaces is a central challenge in computational immunology, with direct implications for vaccine development, cancer immunotherapy, and autoimmune disease. Despite rapid advances in generative protein modeling, there is currently no systematic benchmark evaluating these methods in the highly constrained and immunologically relevant setting of peptide–MHC presentation and TCR recognition. Here, we present two complementary contributions. First, we introduce a multi-stage computational pipeline for peptide design in predefined TCR-pMHC contexts, integrating generative modeling with sequence optimization and structure-based filtering. Second, we establish a benchmark for evaluating generative peptide design methods in TCR-pMHC complexes. Using a curated dataset of high-quality crystal structures deposited after the AlphaFold3 training cutoff, we assess state-of-the-art generative approaches for peptide backbone generation, sequence design, and the enrichment of near-native solutions. We explicitly examine whether different backbone generation strategies respect the geometric constraints of the MHC binding groove and recover native-like peptide conformations. Our results reveal substantial method-dependent differences: some generative strategies fail systematically in the groove-bound peptide setting, whereas others generate physically plausible backbones with varying accuracy and conformational diversity. We further show that enforcing anchor constraints strongly influences peptide conformations at non-anchor positions, highlighting a trade-off between structural accuracy and conformational sampling. To enable fair and reproducible comparison, we introduce a standardized, multi-stage scoring protocol that integrates MHC binding prediction, physics-based energy evaluation, and independent structure prediction confidence metrics to enrich near-native designs from large candidate pools. Together, this work establishes the first comprehensive pipeline and benchmark for generative peptide design at TCR-pMHC interfaces and provides practical guidelines for developing peptide design workflows and evaluating generative models in immunologically constrained protein design settings.

## Introduction

T cell receptors (TCRs) recognize antigenic peptides presented by major histocompatibility complex (MHC) molecules, forming the molecular foundation of adaptive immunity [1]. This trimolecular interaction acts as a central checkpoint of immune decision-making, determining whether immune responses are mounted against pathogens and tumors or, in the case of misrecognition, against self-tissues in autoimmune diseases.

Significant opportunities for clinical intervention revolve around the identification of antigenic peptides. For example, cancer immunotherapy relies on identifying tumor-specific neoantigens that elicit robust T cell responses [2]. These neoantigens can then be targeted using adoptive T cell therapies with engineered TCRs [3]. Conversely, in autoimmune diseases, self-peptides presented by specific HLA alleles can trigger abnormal TCR recognition and pathological immune responses; identifying these peptides could inform strategies for tolerance induction and disease modulation [4].

Motivated by these therapeutic applications, substantial effort has been devoted to developing computational tools for peptide discovery and design. However, current approaches struggle to capture the high specificity of TCR recognition and the extensive polymorphism of MHC molecules across human populations, complicating the therapeutic manipulation of TCR–pMHC interactions.

These limitations have shifted attention toward structure-based modeling approaches enabled by recent advances in protein structure prediction. In this domain, AlphaFold2 [5] and more recently AlphaFold3 (AF3) [6] achieved near-experimental accuracy in predicting protein structures from sequence alone. In parallel, RoseTTAFold introduced a three-track neural network architecture that enabled accurate structure prediction with reduced computational requirements [7]. Building on these foundations, generative models have emerged as powerful tools for *de novo* protein design, extending beyond structure prediction to the creation of new protein structures and sequences. RFdiffusion pioneered the application of denoising diffusion probabilistic models to protein backbone generation, enabling the creation of diverse, designable protein structures conditioned on geometric and contextual constraints, including binding partners [8]. Subsequent developments have extended these capabilities. RFdiffusion1 [8] (RFd1) introduced diffusion-based backbone generation with sequence design performed separately. RFdiffusion3 (RFd3) jointly models backbone and sidechain atoms to better integrate sequence and interaction constraints [9]. BindCraft integrates iterative generative design with AlphaFold2-based filtering to optimize binders [10].

Despite recent advances, the TCR–pMHC system poses distinct modeling challenges. Extreme sequence diversity and structural variability across TCRs and peptide ligands lead to heterogeneous binding modes and poor generalization to unseen TCR–epitope pairs [11]. Although MHC-bound peptides adopt constrained, allele-specific conformations [12, 13], they can undergo context-dependent rearrangements upon TCR engagement. These challenges are exacerbated by the scarcity of high-resolution TCR–pMHC binding data, particularly for rare HLA alleles and non-viral peptides [14]. In addition, the intrinsic flexibility of TCR complementarity-determining region (CDR) loops and peptide backbones complicates docking and interface prediction [15] . Together, these factors make accurate modeling of TCR–pMHC interactions substantially more difficult than standard protein design tasks.

In this work, we focus on peptide discovery, a rapidly growing area with direct relevance to cancer immunotherapy and autoimmune disease. Despite its therapeutic importance and the rapid advancement of generative modeling techniques, computational approaches for peptide design in TCR–pMHC systems remain underdeveloped, and systematic benchmarks for evaluating generative methods are largely lacking. Existing computational models for TCR–pMHC interactions primarily focus on structure prediction [16, 17] and are therefore not readily adaptable to peptide discovery and design tasks.

Our contributions are twofold. First, we introduce a multi-stage computational pipeline for peptide design in predefined TCR–pMHC contexts. Second, we establish a rigorous benchmark for evaluating generative peptide design methods in this highly constrained setting. Our benchmark systematically evaluates de novo peptide reconstruction using RFd1, RFd3, and BindCraft, with ProteinMPNN [18] for sequence design. We curate a dataset of 10 high-quality TCR–pMHC crystal structures deposited after the AlphaFold3 training cutoff, selected using a multi-metric crystallographic quality scoring scheme. Using this dataset, we compare peptide backbone generation and sequence design performance across diverse MHC alleles and peptide lengths.

To enable fair comparison, we introduce a standardized, multi-stage scoring framework that combines MHC binding prediction, physics-based energy evaluation, and structure prediction confidence metrics to enrich near-native solutions from large candidate pools. Finally, we analyze how anchor constraints influence conformational diversity at non-anchor positions, revealing trade-offs between structural accuracy and sampling diversity. Together, these contributions establish a foundation for benchmarking generative models in immunologically relevant protein interfaces and provide practical guidelines for computational design of peptide-based immunotherapy.

## Methods

### De novo peptide design workflow

The proposed *de novo* peptide design workflow is a multi-stage computational pipeline engineered to identify high-affinity peptides for specific TCR-MHC targets (Figure 1). The process integrates generative deep learning for structure and sequence generation, bioinformatic tools for binding prediction, physics-based methods for binding free energy refinement, and state-of-the-art structure prediction for validation. Through progressive filtering and iterative optimization, the pipeline narrows an initial pool of hundreds of thousands of candidates to a tractable number of experimentally viable leads. The workflow comprises four sequential phases, each applying increasingly stringent selection criteria to the outputs of the previous stage.

**Figure 1.**
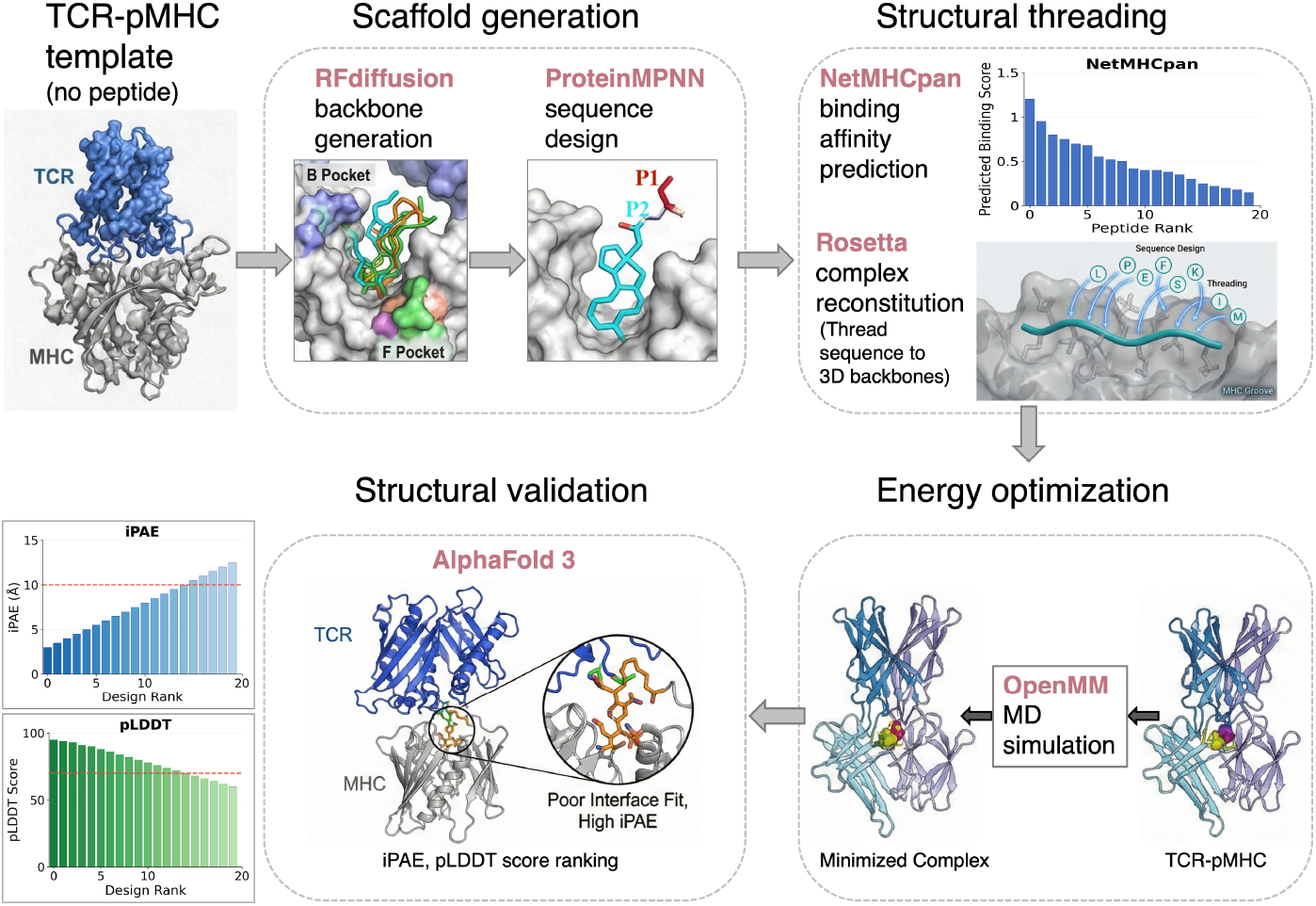
De novo peptide design workflow for high-affinity TCR-MHC targeting. The pipeline comprises four phases: (I) scaffold generation with RFdiffusion and sequence sampling with ProteinMPNN; (II) binding affinity filtering with NetMHCpan and structural threading with Rosetta; (III) energy optimization with OpenMM molecular dynamic (MD) simulation and Rosetta interface analysis, and dual approach using FoldX with machine learning prediction model; (IV) structural validation with AlphaFold3, producing the top 100 designs ranked by interface predicted Local Distance Difference Test (pLDDT).

### Phase I: scaffold generation and sequence sampling

#### Template preparation

To enable controlled evaluation, we first construct a ground-truth dataset of experimentally resolved TCR–pMHC complexes. Namely, we collect high-quality TCR–pMHC crystal structures deposited in the Protein Data Bank (PDB) ^1^ [19] after November 2021 (the AF3 training data cutoff), ensuring that evaluation is not biased by structures seen during AF3 training. From this set, we select 10 template structures that span diverse TCRs and MHC class I contexts (Table 1). We focus on MHC class I complexes because they are more abundantly represented in high-resolution structural data, present shorter peptides (typically 8–11 residues) with well-defined anchor constraints, and exhibit relatively conserved binding geometries across alleles. In contrast, MHC class II molecules bind longer, more variable peptides in open-ended grooves, resulting in greater heterogeneity in binding modes and conformational flexibility, which we leave for future work.

**Table 1.**
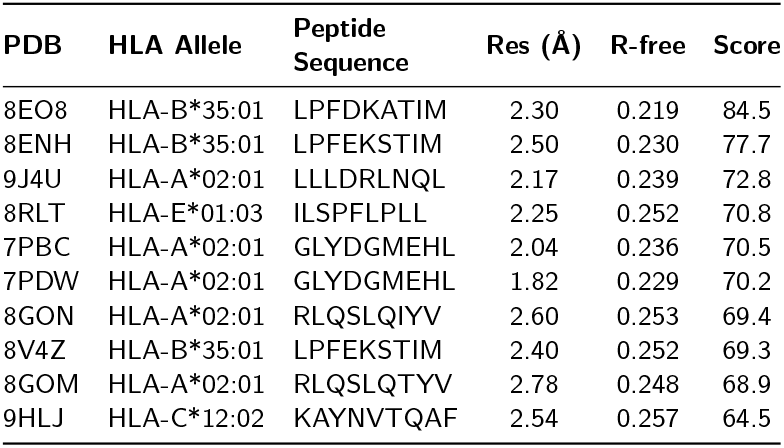
The crystallographic quality metrics, HLA allele assignments, and peptide information for the selected 10 complex templates. Structures are ranked by composite quality (see *Dataset selection and quality assessment*).

For each crystal structure, the native peptide is removed to create a design template that preserves the experimentally determined TCR–MHC interface geometry while treating the peptide as the design target. All atomic coordinates of the TCR *α* and *β* chains, the MHC heavy chain (and *β*_2_-microglobulin for class I), and other non-peptide components are retained, while crystallographic water molecules are removed. The resulting template provides a fixed structural context for all subsequent generative and scoring steps.

#### Backbone generation with RFdiffusion (RFd)

We initially evaluated several generative approaches for peptide backbone generation, including BindCraft [10] and the RFdiffusion family (RFd1-RFd3). Preliminary analyses revealed that BindCraft, which was designed for *de novo* binder generation against exposed protein surfaces, consistently placed generated peptides outside the spatially constrained MHC binding groove rather than within it (Supplementary Figure S1). RFdiffusion2 (RFd2) requires specification of an origin (ORI) token that defines the center of mass for the designed region, a constraint intended for enzyme active site scaffolding rather than groove-filling peptide design. Furthermore, RFd2 documentation explicitly notes that the method is optimized for enzyme scaffolding and recommends the original RFd for binder design applications. Given these limitations, we focused our benchmark on RFd1 [8] and RFd3 [9], both of which support hotspot-based constraints suitable for directing peptide generation within the MHC groove.

Peptide backbone generation is performed using RFd1 [8] and RFd3 [9], denoising diffusion probabilistic models built on RoseTTAFold [20] representations and trained on protein structural data derived from the PDB. These models learn to reverse a noise process applied to known protein structures, enabling the generation of diverse backbones under user-specified structural constraints. Here, RFd1 and RFd3 are configured to generate peptide backbones within the MHC binding groove by enforcing geometric constraints from the surrounding TCR and MHC structures. Correct peptide placement is ensured by specifying *hotspot* residues at canonical anchor positions, which for MHC class I targets correspond to P2 and PΩ (the C-terminal residue) engaging the B and F pockets that determine allele-specific binding [21]. Hotspot constraints guide diffusion sampling by enforcing appropriate distances and orientations between peptide anchors and MHC pocket-lining residues.

For each template structure, we generate 800 peptide backbones of length 8 residues using 200 diffusion steps with a noise schedule optimized for constrained sampling. We design 8-mer backbones because RFd generates residues only between anchor positions; the terminal residue in native complexes outside the anchor span is excluded. The generated backbones are subsequently filtered to remove conformations exhibiting severe steric clashes with the fixed TCR–MHC context.

#### Sequence design with ProteinMPNN

The 800 generated backbones serve as templates for sequence design using ProteinMPNN [18], a graph neural network trained to predict amino acid sequences compatible with specified backbone geometries. ProteinMPNN employs a message-passing architecture that propagates structural information through the protein graph, enabling context-aware sequence predictions that account for both local backbone geometry and longer-range interactions with the surrounding environment. For each backbone template, we configure ProteinMPNN to design sequences for the peptide positions while holding the TCR and MHC sequences fixed at their native identities. We use a sampling temperature of 0.1 to favor high-confidence predictions while maintaining sufficient diversity to explore sequence space. For each backbone, we sample 20 sequences, yielding a total of 16,000 candidate peptide sequences across the 800 backbones. This strategy ensures broad coverage of sequence space compatible with each backbone geometry and increases the likelihood of identifying optimal candidates for subsequent filtering.

#### Sequence completion for MHC class I binding

RFd generation is restricted to the anchor-constrained core of the peptide (positions P2 through PΩ). However, MHC class I molecules require peptides with defined N-terminal residues for proper engagement of the binding groove. To satisfy the 9-mer length typical of most class I alleles, we therefore prepend a P1 residue to each sequence following the ProteinMPNN design step. P1 residues are introduced in a structure-aware manner, accounting for the geometry of the A pocket at the N-terminus of the MHC binding groove. No allele-specific preferences are imposed at this position; instead, each of the 20 amino acids is considered individually at P1. This sequence completion step expands the 320,000 anchor-constrained core sequences into full-length peptides suitable for downstream MHC binding prediction.

### Phase II: Binding affinity filtering and structural threading

With backbone and sequence generation complete, we next filter candidates based on predicted MHC binding affinity and prepare full-atom structural models for physics-based evaluation.

#### MHC binding affinity prediction with NetMHCpan

We apply an initial filtering step to the 320,000 designed sequences based on predicted MHC binding affinity using NetMHCpan-4.1 [22]. NetMHCpan employs a neural network trained on experimentally measured binding affinities and mass spectrometry–identified MHC ligands to predict peptide–MHC binding across a wide range of human HLA alleles. The model outputs both a predicted binding affinity (IC_50_ in nM) and a percentile rank score indicating how a given peptide compares to a reference set of random peptides. For each designed sequence, binding affinity is predicted for the specific HLA allele present in the corresponding template structure. Sequences are ranked by predicted binding strength, and the top 2,000 candidates (representing 0.625% of all designs) are selected for further processing. This stringent filtering step removes sequences with unfavorable anchor residues, incompatible charge distributions, or other features that would preclude stable MHC binding. The resulting 2,000 candidates retain sufficient sequence diversity for downstream optimization while substantially reducing the computational cost of subsequent structure-based analyses.

#### Complex reconstitution with Rosetta SimpleThreadingMover

The selected 2,000 sequences are integrated with their corresponding backbone structures to enable physics-based evaluation. This reconstitution is performed using the SimpleThreadingMover within the Rosetta macromolecular modeling suite [20], which threads a specified amino acid sequence onto an existing backbone template while optimizing side-chain conformations.

For each of the 2,000 selected sequences, the SimpleThreadingM identifies the backbone from which the sequence was derived and threads the designed sequence onto this template. This process replaces the placeholder residues in the backbone with the designed amino acids, followed by side-chain rotamer optimization using Rosetta’s Packer algorithm. The Packer explores discrete rotamer states for each residue, selecting conformations that minimize steric clashes and maximize favorable interactions. Threading and optimization are performed in the context of the full TCR–pMHC complex, with TCR and MHC side chains held fixed to preserve the experimentally determined interface geometry.

The output of this step is a set of 2,000 full-atom models of designed TCR–peptide–MHC complexes, which are then used for subsequent energy evaluation.

### Phase III: Dual-track energy optimization

The 2,000 threaded complexes undergo energy optimization to ensure physical feasibility and identify the most stable designs. This optimization employs two parallel scoring tracks that capture different aspects of protein stability and binding energetics. Track A employs FoldX force field calculations [23] combined with a machine learning classifier, while Track B uses OpenMM [24] for molecular dynamics minimization, followed by Rosetta [20] interface analysis. The two tracks provide independent assessments that are combined through consensus ranking to identify the most promising candidates.

#### FoldX force field with TRAIT machine learning

The first scoring track applies the FoldX empirical force field [23] to compute comprehensive energy profiles for each TCR-pMHC complex. FoldX calculations yield three categories of energy terms that characterize different aspects of complex stability and binding.

**Stability energies** quantify the overall structural integrity of the complex through 23 individual terms including van der Waals interactions, backbone and sidechain hydrogen bonds, electrostatics, solvation contributions (polar and hydrophobic), entropy terms for main chain and side chains, and specialized terms for disulfide bonds, cis-peptide bonds, and torsional strain. The RepairPDB function [23] first optimizes side-chain conformations and removes steric clashes before computing these terms. **Individual chain energies** assess the internal stability of each chain in isolation, with particular focus on the peptide chain. These terms identify designs where the peptide sequence is intrinsically unstable independent of the binding context. **Interaction energies** capture the pairwise energetic contributions between chains at the binding interfaces. For TCR-pMHC complexes, we compute interaction energies between TCR*α*-peptide, TCR*β*-peptide, MHC–peptide, and the combined TCR-peptide interfaces. Key terms include interface van der Waals clashes, electrostatic contributions, hydrogen bonding across the interface, and the number of interface residues with steric conflicts.

The FoldX energy terms serve as input features for TRAIT (TCR Recognition of Antigen through Interaction Testing) [25], a logistic regression classifier trained to discriminate favorable from unfavorable TCR-pMHC interactions. TRAIT employs L1-regularized (Lasso) logistic regression with bootstrap stability analysis to identify the most predictive energy features. Through 100 bootstrap iterations with 5-fold cross-validation, 8 stable features (selection frequency ≥70%) were identified as the core predictive set. The TRAIT classifier outputs a probability score for each design, with higher scores indicating greater predicted likelihood of favorable TCR-pMHC interaction. Designs are ranked by this probability score to produce FoldX force field rankings.

#### OpenMM minimization with Rosetta interface analysis

The second scoring track employs molecular mechanics minimization followed by physics-based interface analysis, providing a complementary assessment grounded in explicit atomic interactions.

**OpenMM energy minimization** uses the Amber14 all-atom force field [24] to optimize the geometry of each TCR-pMHC complex. The minimization protocol begins by adding missing hydrogen atoms to the structure using OpenMM’s Modeller class, ensuring proper protonation states at physiological pH. The system is constructed without explicit solvent (vacuum minimization) using the NoCutoff nonbonded method with HBonds constraints on hydrogen-containing bonds. Energy minimization employs the L-BFGS algorithm through the Langevin integrator framework, running for up to 1,000 iterations or until convergence. The OpenMM minimization is performed with H100 GPU acceleration, enabling rapid processing of the 2,000 candidate structures. For structures that fail initial minimization due to severe clashes or missing atoms, the PDBFixer utility is applied to repair the structure before a second minimization attempt. The minimized structures and their final potential energies are recorded for subsequent analysis.

**Rosetta interface analysis** applies the InterfaceAnalyzerMover from the Rosetta macromolecular modeling suite [20] to quantify binding energy at the TCR-pMHC and peptide-MHC interface. For each minimized structure, interface analysis is performed with two partner definitions: 1) quantifies the binding for TCR-pMHC interface and yields metrics interface Δ*G*, separated interface energy (dG), and interface buried surface area (ΔSASA); 2) quantifies peptide binding within the MHC groove and yields Δ*G*, dG and (ΔSASA). Additionally, explicit binding energy calculations decompose the complex energy: Δ*E*_binding_ = *E*_complex_ − (*E*_TCR_ + *E*_pMHC_), where each component is scored using Rosetta’s energy function after splitting the complex into constituent parts. The energy score rankings are generated based on a composite score combining the OpenMM minimized energy, interface Δ*G* values, and Rosetta binding energy, with appropriate weighting to balance the contributions of each metric.

#### Consensus ranking and selection

The independent rankings from FoldX with TRAIT and OpenMM with Rosetta are combined through consensus ranking to identify designs that perform well across both scoring paradigms. For each design, the geometric mean of its ranks in both tracks is computed: 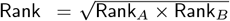.

This consensus approach prioritizes designs that achieve favorable scores under both the machine learning-based assessment (which captures statistical patterns learned from experimental data) and the physics-based assessment (which evaluates explicit atomic interactions). Designs ranking highly in one track but poorly in the other receive intermediate consensus ranks, while designs performing well in both tracks are promoted to top positions. Based on the consensus ranking, the top 100 designs are selected for high-fidelity structural validation. This selection threshold retains approximately 5% of the threaded structures, representing designs that have demonstrated favorable energetics under both scoring paradigms.

### Phase IV: structural validation with AlphaFold3

The final phase employs an independent structure prediction method to validate the plausibility of top-ranked designs. The 100 consensus-selected designs undergo structural validation using AlphaFold3 (AF3) [6], which serves as an independent assessment for structural plausibility of the designed TCR-pMHC complexes. AF3 extends the capabilities of AlphaFold2 [5] to protein complexes, providing confidence metrics that reflect the reliability of predicted inter-chain contacts. For each designed complex, we extract the peptide sequence and predict the structure of the peptide-MHC complex using AF3. The key validation metric is the interface predicted Local Distance Difference Test (interface pLDDT), calculated as the mean pLDDT of peptide atoms within 5 Å of any MHC or TCR atom. Interface pLDDT scores above 70 indicate high-confidence local structure predictions where the peptide is predicted to adopt a well-defined conformation with reliable atomic positions at the binding interface. Application of the interface pLDDT ≥ 70 threshold to the 1000 consensus-ranked designs yields approximately 100 validated complexes. These designs represent the highest-confidence candidates from the initial 320,000 sequences, having passed filters for MHC binding affinity, dual-track energy assessment, and structural plausibility. The top 100 designs ranked by interface pLDDT are selected for detailed analysis, representing candidates with the most reliable predicted interface geometries.

### Dataset selection and quality assessment

The assembly of a high-quality benchmark dataset required consideration of both structural quality and the potential for data leakage from structure prediction models’ training sets. We queried the PDB using a combination of search terms with the boolean expression “(T cell receptor OR TCR) AND (HLA OR MHC) AND peptide” with the experimental method restricted to X-ray diffraction after AF3 release date. We selected the final 10 benchmark templates using a composite crystallographic quality score. This score incorporates seven metrics: resolution, R-free, R-work, R-gap, I/*σ*, CC_1*/*2_, and completeness. We performed Principal Component Analysis (PCA) on these metrics following established practices for index construction [26]. Metrics where lower values indicate better quality (resolution, R-free, R-work, R-gap) were inverted prior to analysis, and all metrics were standardized using z-score normalization. Weights were derived using variance-weighted loadings: 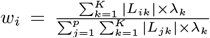, where *L*_*ik*_ is the loading of metric *i* on component *k*, and *λ*_*k*_ is the variance explained by component *k*.

The first six principal components (PC) explained 100% of the total variance: PC1 (40.3%) captured resolution-related quality, PC2 (17.4%) reflected R-factor reliability, PC3 (14.6%) represented data completeness and signal strength, and PC4 (11.6%) indicated overfitting. The PCA-derived weights were: R-work (16%), resolution (15%), R-gap (15%), I/*σ* (15%), R-free (14%), completeness (14%), and CC_1*/*2_ (12%). These derived weights validated R-work as the primary determinant of structure quality [27]. Resolution-specific R-factor thresholds were applied, ranging from R-free ≤ 0.18 at near-atomic resolution (≤1.8 Å) to R-free ≤ 0.22 at lower resolution (3.0– 4.0 Å). The 10 highest-scoring structures spanning diverse HLA alleles were selected as benchmark templates (Table 1).

## Results

We benchmarked RFd1 and RFd3 for *de novo* peptide backbone generation in TCR-pMHC complexes. Our evaluation employed a curated dataset of 10 high-quality crystal structures spanning four distinct HLA alleles (HLA-A*02:01, HLA-B*35:01, HLA-C*12:02, and HLA-E*01:03), with 1000 designs generated per method. All generated peptides were subsequently validated using AF3 structure prediction and Rosetta-based interface analysis. AF3’s interface predicted Local Distance Difference Test (I-pLDDT), calculated as the mean pLDDT of peptide atoms within 5Å of any MHC or TCR atom [28], served as a key metric for assessing local structural confidence at the binding interface.

Our analysis reveals a fundamental trade-off between structural fidelity and predicted binding propensity (Table 2). RFd1 consistently outperformed RFd3 on structure-based quality metrics: peptide backbone RMSD was significantly lower (6.84 vs. 7.71 Å; *p <* 10^−8^), interface binding energy more favorable (Δ*G* = −76.5 vs. −60.1 REU; *p <* 10^−80^), and structural confidence higher for both peptide pLDDT (76.7 vs. 72.6) and interface pLDDT (85.1 vs. 83.8). AlphaFold3 validation metrics similarly favored RFd1, with higher iPTM (0.73 vs. 0.71) and reduced peptide–TCR*β* alignment error (PAE = 7.13 vs. 8.65 Å). Notably, RFd1 designs exhibited lower variance in interface energy, indicating more consistent generation of physically plausible conformations.

**Table 2.** Overall comparison of AF3 evaluation metrics between RFd1 and RFd3 for de novo peptide design in TCR-pMHC complexes. Values shown as mean *±* std. Bold indicates significantly better performance (*p <* 0.05, Mann-Whitney U test).

| Metric | RFd1 (n=1000) | RFd3 (n=1000) | p-value |
| --- | --- | --- | --- |
| iPTM | **0.73 ± 0.10** | 0.71 ± 0.09 | $1.7 \times 10^{-6}$ |
| pTM | **0.78 ± 0.07** | 0.77 ± 0.07 | $4.9 \times 10^{-6}$ |
| AF3 Ranking Score | **0.75 ± 0.10** | 0.73 ± 0.09 | $1.3 \times 10^{-6}$ |
| Peptide RMSD (Å) | **6.84 ± 3.47** | 7.71 ± 3.38 | $7.5 \times 10^{-9}$ |
| PAE<sub>pep-TCRβ</sub> (Å) | **7.13 ± 4.72** | 8.65 ± 4.93 | $8.5 \times 10^{-20}$ |
| $\Delta G_{\text{interface}}$ (REU) | **-76.5 ± 21.7** | -60.1 ± 44.5 | $2.3 \times 10^{-88}$ |
| peptide pLDDT | **76.7 ± 10.6** | 72.6 ± 10.6 | $1.6 \times 10^{-20}$ |
| interface pLDDT | **85.1 ± 4.4** | 83.8 ± 3.6 | $2.1 \times 10^{-12}$ |
| $P_{\text{binder}}$ | 0.52 ± 0.10 | **0.60 ± 0.09** | $2.1 \times 10^{-72}$ |

In contrast, RFd3 achieved significantly higher predicted binder probability (*P*_binder_ = 0.60 vs. 0.52; *p <* 10^−70^). This metric, derived from a machine learning classifier TRAIT trained on experimental binding data, captures features predictive of productive TCR-peptide interactions. The divergence between structural accuracy and binding propensity suggests that RFd3 explores a conformational landscape distinct from native geometries yet enriched for binding-competent features.

Template-specific analysis revealed substantial heterogeneity in method performance across TCR–pMHC contexts (Table 3). RFd1 demonstrated clear advantages for HLA-B*35:01 complexes (PDBs 8EO8, 8ENH, 8V4Z) and HLA-E*01:03 (8RLT), achieving iPTM values of 0.79–0.82 and favorable interface energy. However, for select templates, RFd3 showed competitive or superior structural metrics: in PDB 8GON (HLA-A*02:01), RFd3 achieved higher iPTM (0.74 vs. 0.61) and peptide pLDDT (80.0 vs. 73.0); in PDB 9HLJ (HLA-C*12:02), RFd3 showed lower PAE_*β*_ (5.79 vs. 6.82 Å). These context-dependent patterns suggest that method performance may vary with allele-specific groove geometries and native peptide conformational flexibility. Noted, RFd3 achieved higher *P*_binder_ across all ten templates without exception, underscoring its consistent advantage in predicted binding propensity.

**Table 3.** Per-PDB comparison of key AF3 metrics. Bold indicates the better method for each PDB-metric pair. Each PDB has 100 designs per method. Abbreviations: RMSD = peptide RMSD, PAE_*β*_ = PAE_pep-TCR*β*_, Δ*G* = interface energy, pLDDT_pep_/pLDDT_int_ = peptide/interface pLDDT, *P*_bind_ = binder probability.

| PDB | Method | iPTM | RMSD (Å) | PAE$_{\beta}$ (Å) | $\Delta G$ | pLDDT<sub>pep</sub> | pLDDT<sub>int</sub> | $P_{\text{bind}}$ |
| --- | --- | --- | --- | --- | --- | --- | --- | --- |
| 7PBC | RFd1 | **0.81** | **4.11** | **4.38** | **-81.1** | **80.8** | **88.8** | 0.39 |
|  | RFd3 | 0.73 | 5.42 | 7.80 | -52.6 | 65.7 | 84.0 | **0.48** |
| 7PDW | RFd1 | **0.82** | 8.88 | **3.81** | **-84.2** | **84.1** | **89.7** | 0.39 |
|  | RFd3 | 0.78 | **8.47** | 5.46 | -51.5 | 74.4 | 86.9 | **0.49** |
| 8ENH | RFd1 | **0.79** | **5.49** | **6.09** | **-70.1** | **75.3** | **85.0** | 0.54 |
|  | RFd3 | 0.73 | 6.45 | 8.04 | -65.7 | 66.0 | 81.3 | **0.65** |
| 8EO8 | RFd1 | **0.82** | **5.99** | **4.70** | -59.3 | **79.4** | **86.4** | 0.51 |
|  | RFd3 | 0.74 | 7.21 | 9.02 | **-63.3** | 61.6 | 80.8 | **0.61** |
| 8GOM | RFd1 | **0.58** | 9.64 | **14.77** | **-80.8** | 63.5 | 79.1 | 0.59 |
|  | RFd3 | 0.57 | **9.51** | 18.01 | -38.3 | **78.4** | **82.0** | **0.66** |
| 8GON | RFd1 | 0.61 | 8.43 | 12.53 | **-76.7** | 73.0 | 82.1 | 0.55 |
|  | RFd3 | **0.74** | **8.43** | **7.19** | -73.6 | **80.0** | **86.3** | **0.62** |
| 8RLT | RFd1 | **0.76** | **6.97** | **3.81** | **-85.1** | **84.4** | **88.5** | 0.57 |
|  | RFd3 | 0.70 | 9.07 | 8.59 | -72.3 | 74.8 | 84.4 | **0.68** |
| 8V4Z | RFd1 | 0.79 | 6.82 | 6.35 | **-72.7** | **73.6** | 85.2 | 0.55 |
|  | RFd3 | **0.82** | **6.79** | **5.41** | -59.6 | 72.1 | **85.5** | **0.64** |
| 9HLJ | RFd1 | 0.69 | **2.92** | 6.82 | -71.2 | 75.3 | 83.0 | 0.63 |
|  | RFd3 | **0.72** | 7.18 | **5.79** | **-72.3** | **80.2** | **85.7** | **0.67** |
| 9J4U | RFd1 | **0.65** | 9.14 | **8.02** | **-83.6** | **77.2** | **83.2** | 0.46 |
|  | RFd3 | 0.61 | **8.61** | 11.19 | -52.0 | 72.4 | 81.3 | **0.50** |

To investigate the role of geometric constraints in guiding peptide generation, we evaluated eight hotspot configurations ranging from MHC-only constraints (4-12 residues) to combined MHC+TCR constraints incorporating complementarity-determining region (CDR) loop contacts (8-20 total residues; Table 4). Contrary to expectation, increasing constraint density did not monotonically improve backbone quality. RFd1 achieved favorable metrics across 7 of 8 configurations, with optimal performance at moderate constraint levels: the lowest RMSD (6.28 Å) occurred with 16 hotspots, while the most favorable interface energy (−83.8 REU) was observed with 12 MHC-only hotspots. RFd3 exhibited greater sensitivity to constraint configuration, benefiting from combined MHC+TCR constraints that may better guide its all-atom sampling. These findings highlight constraint optimization as an important tunable parameter, with moderate constraint sets (8–12 hotspots) generally balancing geometric accuracy against conformational diversity.

**Table 4.** Per-experiment comparison between RFd1 and RFd3 across eight hotspot configurations. Bold indicates better performance. Hotspot (HS) counts shown in parentheses represent total constraint residues. MHC-only experiments (EXP1-3) constrain peptide-MHC groove interactions using 4, 8, or 12 MHC residues flanking anchor positions. Combined MHC+TCR experiments (EXP4-8) additionally constrain peptide-TCR contacts using CDR loop residues from both TCR*α* and TCR*β* chains.

| Experiment | Method | N | iPTM | RMSD | PAE<sub>β</sub> | $\Delta G$ | pLDDT<sub>prep</sub> | pLDDT<sub>int</sub> | $P_{\text{bind}}$ |
| --- | --- | --- | --- | --- | --- | --- | --- | --- | --- |
| EXP1 (4 HS) | RFd1 | 140 | **0.738** | **6.63** | **6.92** | **-78.6** | **79.0** | **86.0** | **0.50** |
|  | RFd3 | 74 | 0.700 | 8.04 | 10.23 | -66.7 | 75.5 | 83.9 | **0.61** |
| EXP2 (8 HS) | RFd1 | 135 | 0.725 | **7.80** | 7.99 | **-78.3** | **75.5** | **84.8** | 0.51 |
|  | RFd3 | 111 | **0.735** | 8.73 | **7.68** | -58.8 | 70.7 | 84.1 | **0.59** |
| EXP3 (12 HS) | RFd1 | 193 | **0.735** | **6.49** | **6.48** | **-83.8** | **79.4** | **86.0** | 0.47 |
|  | RFd3 | 93 | 0.700 | 7.32 | 8.69 | -50.3 | 72.7 | 83.7 | **0.58** |
| EXP4 (8 HS) | RFd1 | 100 | **0.759** | **6.86** | **5.85** | **-74.9** | **78.1** | **85.7** | 0.51 |
|  | RFd3 | 198 | 0.700 | 8.04 | 9.20 | -51.8 | 70.7 | 83.0 | **0.59** |
| EXP5 (14 HS) | RFd1 | 120 | **0.708** | **7.24** | **7.51** | **-76.4** | **75.7** | 84.4 | 0.52 |
|  | RFd3 | 98 | 0.703 | 8.35 | 9.46 | -67.3 | 74.5 | **84.4** | **0.58** |
| EXP6 (16 HS) | RFd1 | 112 | 0.729 | **6.28** | 7.90 | -66.0 | 73.8 | 84.1 | 0.58 |
|  | RFd3 | 148 | **0.736** | 7.43 | **7.80** | **-72.6** | **77.2** | **85.4** | **0.62** |
| EXP7 (14 HS) | RFd1 | 68 | **0.751** | 7.29 | **6.88** | **-71.1** | **74.4** | **84.7** | 0.54 |
|  | RFd3 | 169 | 0.723 | **7.29** | 8.05 | -55.1 | 69.4 | 83.3 | **0.61** |
| EXP8 (20 HS) | RFd1 | 132 | **0.723** | **6.46** | **7.51** | **-74.3** | **74.7** | **84.4** | 0.55 |
|  | RFd3 | 109 | 0.705 | 6.67 | 8.88 | -64.9 | 72.4 | 83.3 | **0.61** |

To illustrate these findings, we examined PDB 8GON (HLA-A*02:01) as a representative case study (Figure 2). Structural superposition of the native 9-mer peptide (Figure 2a) with an RFd3-generated 8-mer (Figure 2b) reveals that while anchor positions engaging the B and F pockets of the MHC groove are largely preserved, the central peptide region exhibits notable conformational divergence. This observation is consistent with the design strategy. The anchor constraints enforce groove engagement, but the solvent-exposed central residues retain conformational freedom during diffusion sampling.

**Figure 2.**
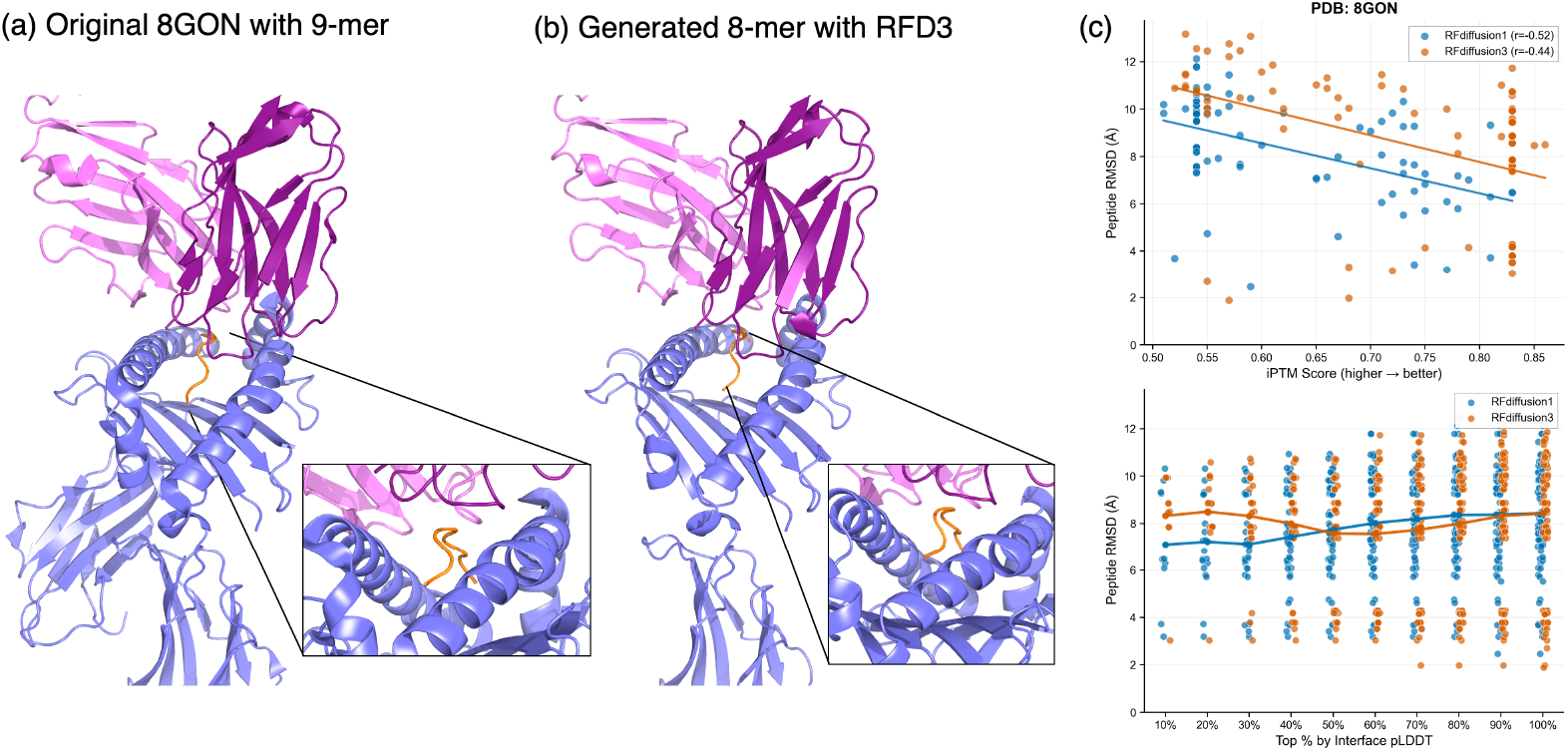
Structural and quantitative comparison of native and generated peptide conformations in TCR-pMHC complexes. **(a)** Native TCR–pMHC crystal structure (PDB: 8GON, HLA-A*02:01) with the 9-mer peptide (orange) bound in the MHC groove (blue); TCR *α* and *β* chains shown in pink and magenta, with magnified view of peptide within the binding groove. **(b)** Representative RFd3-generated 8-mer peptide threaded onto the same TCR-MHC scaffold, showing small differences in the central peptide region. **(c)** Correlation between structural quality metrics and peptide backbone RMSD for PDB 8GON. Upper panel: iPTM score versus RMSD shows negative correlation for both RFd1 and RFd3. Lower panel: RMSD distribution across interface pLDDT percentile bins.

Quantitative analysis of the relationship between confidence metrics and structural accuracy provides insight into the utility of AlphaFold3-derived scores for design filtering (Figure 2c). For PDB 8GON, peptide backbone RMSD showed moderate negative correlation with iPTM score for both RFd1 (*r* = −0.52) and RFd3 (*r* = −0.44), indicating that higher-confidence predictions tend to adopt more native-like conformations. RFd1 achieved lower RMSD values across the iPTM range, consistent with its superior structural fidelity observed in aggregate metrics. When designs were stratified by interface pLDDT percentile, RFd1 maintained consistently lower RMSD than RFd3 across top 40% confidence thresholds. Both methods showed only modest enrichment for native-like conformations at higher pLDDT cutoffs. These results suggest that while iPTM and interface pLDDT provide useful ranking criteria, they capture complementary aspects of structural quality and should be combined with physics-based metrics for optimal design selection.

The multi-stage pipeline demonstrated effective enrichment of high-quality designs from large candidate pools. Shown in Table 5, proposed workflow was starting from 800 backbone conformations per template, ProteinMPNN sequence sampling were expanded the search space to 320,000 candidates. Sequential filtering through NetMHCpan binding prediction, dual-track energy optimization, and AlphaFold3 validation (pLDDT_int_ ≥ 70) yielded 100 validated designs. Method-specific attrition patterns emerged: RFd1-generated backbones achieved higher pass rates through structural validation, consistent with their superior geometric quality metrics. However, among designs passing all filtering stages, RFd3 candidates exhibited comparable or superior predicted binding propensity.

**Table 5.**
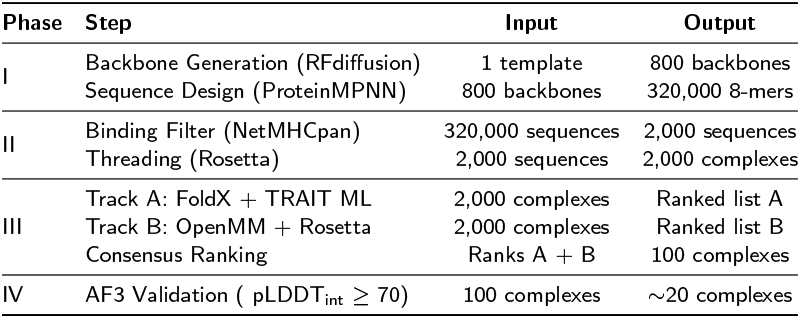
Four-phase computational pipeline for de novo peptide design. Starting from a single TCR–pMHC template, the pipeline generates 320,000 candidate sequences, filters by predicted MHC binding, ranks using consensus scoring from two complementary tracks (energy-based and machine learning-based), and validates with AlphaFold3.

These findings carry practical implications for peptide design workflows. For applications requiring faithful recapitulation of native geometries, RFd1’s superior structural fidelity is advantageous for structure-guided optimization of known epitopes or scaffold transplantation. For exploratory campaigns seeking novel peptides with high binding potential, RFd3’s broader conformational sampling may identify candidates inaccessible to geometry-constrained approaches. An ensemble strategy combining outputs from both methods, followed by stringent multi-stage filtering, may yield the most diverse and high-quality candidate pools for downstream experimental validation.

## Conclusions

Here, we present a multi-stage computational framework for peptide design in TCR–pMHC systems together with a benchmark for evaluating generative peptide design methods. We propose a four-phase pipeline designed to balance diversity, physical realism, and computational efficiency in peptide design. Phase I emphasizes broad exploration of sequence and conformational space via large-scale generative sampling, whereas subsequent phases apply progressively more stringent filters based on binding prediction (Phase II), energetic evaluation (Phase III), and structure prediction validation (Phase IV). This staged design allows inexpensive methods to eliminate low-quality candidates early, reserving computationally intensive analyses for the most promising designs.

Our pipeline integrates complementary methods at each stage, such as sequence-based versus structure-based and learning-based versus physics-based approaches, thereby reducing method-specific biases and achieving more robust performance than any single approach. For instance, our pipeline integrates generative backbone modeling and sequence design using models with different architectures and inductive biases: RFdiffusion generates structures via a diffusion process conditioned on structural context, whereas ProteinMPNN uses a graph neural network to assign sequences to a given backbone. This diversity in modeling assumptions reduces reliance on any single approach and helps mitigate correlated failure modes. Furthermore, sequence-based MHC binding prediction with NetMHCpan provides complementary information about peptide–MHC compatibility that does not depend on conformational energy. We then assess peptide stability and binding energetics using two independent scoring tracks: one that combines empirical force-field evaluation with learned statistical patterns (FoldX and TRAIT), and another that applies molecular mechanics minimization followed by physics-based interface analysis (OpenMM and Rosetta). By integrating these independent assessments through consensus ranking, the pipeline better identifies high-quality candidates than any individual scoring method.

Our results suggest that peptide design benefits from an explicit refinement stage in addition to the initial generative sampling. After identifying designs that pass stringent structural validation, localized refinement further improves peptide conformations while preserving the overall TCR–pMHC structural context. Our refinement focuses on parts of the peptide that were added after initial generation, such as the P1 position, because those regions are more uncertain and benefit from additional structural adjustment. Consistent gains in structural plausibility and predicted MHC binding affinity support the conclusion that generative modeling and targeted refinement serve complementary roles. Together, these findings suggest that iterative, structure-aware refinement is a valuable strategy for enhancing high-confidence peptide designs and can justify the additional computational cost in focused design workflows.

A limitation of this study is that we benchmark peptide design in an idealized setting where the TCR and MHC conformations are known from crystallography. In most clinical applications, however, only the TCR and MHC sequences are available. While structure prediction models can infer geometries from sequence, this inference introduces uncertainty, particularly in flexible regions such as TCR CDR loops and, in some cases, peptide-induced deformation of the MHC binding groove. Hence, fully *de novo* peptide discovery requires explicitly accounting for conformational variability through flexible docking and loop remodeling, as well as evaluating whether the designed sequences adopt the predicted structural conformations and lead to immunologically active complexes.

In addition, our benchmark is limited to a small set of MHC class I alleles from a carefully curated set of 10 high-quality templates, and therefore does not capture the full diversity of TCR–pMHC interactions. Extending this framework to MHC class II will require addressing additional layers of complexity, as class II molecules present longer and more variable peptides in an open-ended binding groove, giving rise to register ambiguity, where the same peptide can bind in multiple overlapping alignments, as well as increased heterogeneity in peptide conformations and binding modes. In future work, we will expand allele coverage and extend the benchmark to MHC class II molecules as additional high-resolution structures become available, enabling broader and more comprehensive peptide discovery.

## Data availability

The data underlying this article are available in RCSB PDB at http://rcsb.org, and can be accessed with 7PBC, 7PDW, 8EO8, 8ENH, 8GOM, 8GON, 8RLT, 8V4Z, 9HLJ and 9J4U. The application code for the software and a step-by-step tutorial are available at https://github.com/Yale-CompBio/pepbench-tcr.

