## Supplemental Figure S1 for "Benchmarking generative scaffold design methods for peptide engineering in TCR–MHC complexes"

Supplementary

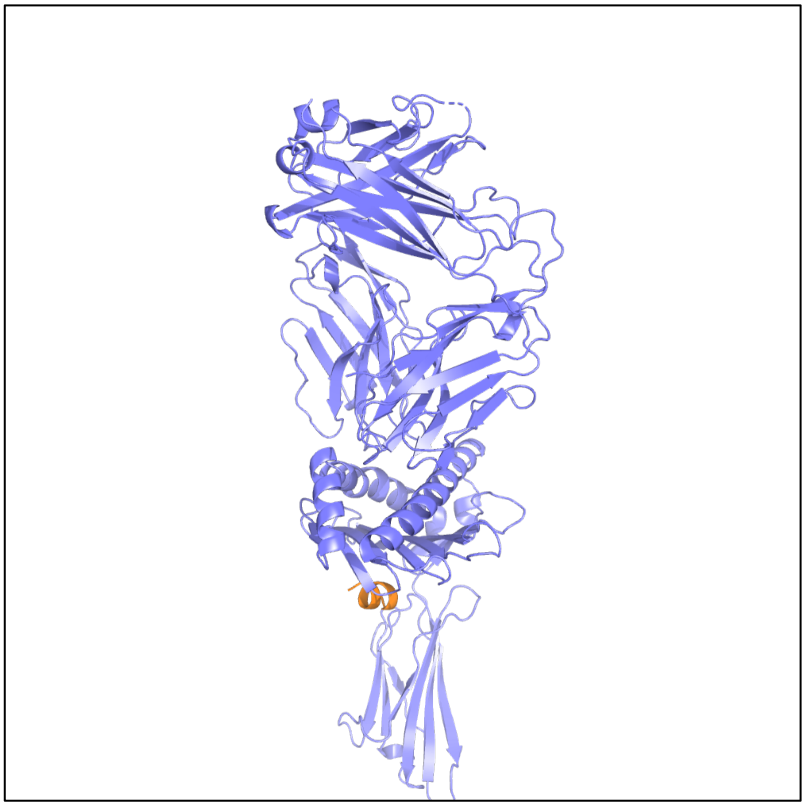

Figure S1: The *de novo* peptide design by BindCraft generated a peptide outside the MHC groove.
